# A Meta-analysis of 2D vs. 3D Ovarian Cancer Cellular Models

**DOI:** 10.1101/2022.12.05.519144

**Authors:** Rachel Kerslake, Birhanu Belay, Suzana Panfilov, Marcia Hall, Ioannis Kyrou, Harpal S. Randeva, Jari Hyttinen, Emmanouil Karteris, Cristina Sisu

**Author notes:** Correspondence: (CS), (EK).

## Abstract

Three-dimensional (3D) cancer models are revolutionizing research, allowing for the recapitulation of *in vivo* like response through the use of an *in vitro* system, more complex and physiologically relevant than traditional mono-layer culture. Cancers such as ovarian (OvCa), are prone to developing resistance and are often lethal, and stand to benefit greatly from the enhanced modelling emulated by 3D culture. However current models often fall short of predicted response where reproducibility is limited owing to the lack of standardized methodology and established protocols. This meta-analysis aims to assess the current scope of 3D OvCa models and the differences in genetic profile presented by a vast array of 3D cultures. A meta-analysis of the literature (Pubmed.gov) spanning 2012 – 2022, was used to identify studies with comparable monolayer (2D) counterparts in addition to RNA sequencing and microarray data. From the data 19 cell lines were found to show differential regulation in their gene expression profiles depending on the bio-scaffold (i.e. agarose, collagen or Matrigel) compared to 2D cell cultures. Top genes differentially expressed 2D vs. 3D include C3, CXCL1, 2 and 8, IL1B, SLP1, FN1, IL6, DDIT4, PI3, LAMC2, CCL20, MMP1, IFI27, CFB, and ANGPTL4. Top Enriched Gene sets for 2D vs. 3D include IFN-α and IFN-γ Response, TNF-α signalling, IL-6-JAK-STAT3 signalling, angiogenesis, hedgehog signalling, apoptosis, epithelial mesenchymal transition, hypoxia, and inflammatory response. Our transversal comparison of numerous scaffolds allowed us to highlight the variability that can be induced by these scaffolds in the transcriptional landscape as well as identifying key genes and biological processes that are hallmarks of cancer cells grown in 3D cultures. Future studies are needed to identify which is the most appropriate in vitro/preclinical model to study tumour microenvironment.

**Summary:** Ovarian cancer is one of the most lethal forms of female cancers. Cell culture is often the go to model to study the molecular processes of cancer. However, this is an oversimplification of the reality. 3D tissue culture has been developed to address the cell culture limitations and to provide a more realistic model of the system studied. Cells grown in 3D represent better the human tumour microenvironment. This meta-analysis is exploring the use of 3D tissue culture as a model of ovarian cancer. Our analysis shows that ovarian cancer cells grown in 3D exhibit enhanced regulation in processes pertinent to tumour development and progression. We identify a panel of genes associated with specific 3D growth conditions that could be used as conditional markers. Finally, we present an overview of the state-of-art of 3D culture with an extensive profile of the genes and pathways enhanced in ovarian cancer models.

## 1. Introduction

### Ovarian Cancer

Ovarian cancer (OvCa) is one of the most lethal gynaecological malignancies of the 21^st^ century. Affecting over 313,000 women worldwide, OvCa typically presents at a late stage with non-specific symptoms, causing a detriment to survival outcomes, which fall as low as 20% [2]. The metabolic processes involved in OvCa aetiology however remain poorly understood. There are three main histological types of OvCa. Epithelial OvCa, accounts for 90% of all cases, with high grade serous ovarian cancer (HGSOC – 70%) being the most prevalent of the five subtypes as well as the most lethal [2]. Other subtypes include low grade serous ovarian cancer (LGSOC – 5%), endometrioid adenocarcinoma of the ovary (EAC – 10%), clear cell carcinoma (CCC – 10%) and mucinous adenocarcinoma (MAC < 3%). The least common are germ line and stromal sex cord tumours which cover 10% of cases [3].

In order gain a better understanding of the events that take place within the tumour microenvironment (TME), a model capable of emulating the *in vivo* milieu is required. The use of conventional monolayer cell culture (two-dimensional; 2D) allows for analysis using a controlled *in vitro* environment to investigate physiological, morphological, and biochemical properties of biological systems [4]. Monolayer culture has served as an integral foundation of biological research since the introduction of immortalised HeLa in 1951 paving the way for thousands of subsequent cell lines [5]. Cell models have since proven invaluable in the modelling of normal physiology and diseases including cancer [6].

Nevertheless, monolayer culture has translational limitations, with differences in gene expression, drug response and cell signalling evident when compared to *in vivo* models [7]. Many processes related to tumorigenesis and metastasis are often over-simplified in monocultures [8]. As a result, monolayer culture often fails to recapitulate the complex microenvironment, diffusion gradients and cellular characteristics associated with *in vivo* systems. Thus, leading to variation from predicted response in animal and computational modelling, as well as clinical testing [7], [9].

As global research efforts strive to answer increasingly complex biological questions, there is a greater need for a representative system capable of physiological emulation. Many studies show that the complexities of tissue organisation, differentiation, and gene expression are demonstrated at higher levels in three-dimensional (3D) cell cultures [10], [11]. This set up allows for cells to be grown in an environment that sustains spatial complexities representative of *in vivo* allowing cells to differentiate and interact in a tissue specific manner [12]. Key differences between monolayer and 3D cultures are summarised in Table 1 [6].

**Table 1.** Differences between 2D and 3D cell culture systems [13].

| 2D - Culture | 3D - Culture |
| --- | --- |
| Cells grown in monolayers – biologically simple | Cells form differentiated aggregates, spheroids, or organoids – biologically complex |
| Gene and protein expression differ from *in vivo* | Expression closely mimics *in vivo* |
| Uniform exposure to chemical stimuli; drugs often appear affective | Nonuniform growth results in toxicity profiles and diffusion gradients closely related to *in vivo* |
| Oxygen diffusion is uniform and higher than many *in vivo* structures; thus, augmenting mitochondrial function and ROS production | Oxygen distribution varies, hypoxic cores are evident; closely mimicking *in vivo* variations of many complexes |
| Long term culture can result in genetic drift with epigenetic and morphological changes evident | Growth is typically short term, minimizing genetic drift |
| Can be cheaper and less complex, therefore easily recapitulated in a lab | Requires additional nutrients and biological scaffolds, and can therefore be more expensive and time consuming |
| Established protocols | Limited established protocols |

Further evidence emphasises the importance of the TME for maintained cancer stemness, exerting a significant effect over gene expression [14]. The integration of an extracellular matrix (ECM) i.e., a scaffold, provides the necessary environment for this 3D cellular growth and differentiation [15]. Scaffolds emulate the tissue-tissue interfaces and chemical gradients required within a living system. Recent advancements include 3D organoid systems capable of sustaining a vast array of tumour models including glioblastoma, colon and lung as well as ovarian [16]–[18].

Epithelial OvCa cells grown in 3D, often present with histological features characteristic of the original tumour *in situ* [19]. 3D epithelial OvCa cell lines also present with reduced proliferative rate thought to be enabled by the synthetic ECM [20]. An enhanced response to external stimuli is also evident within OvCa cultures. Thus far 3D OvCa cultures have proven particularly useful as a model of therapeutic resistance; capturing developed resistance to platinum-based therapeutics similar to *in vivo* OvCa response. The OvCa cell line SKOV-3, for example demonstrates a higher degree of chemoresistance to both cisplatin and paclitaxel when cultured in 3D [21]. Moreover, colorectal and pancreatic cancer cells grown in 3D exhibit differential gene expression that is associated with augmented ATP production within 3D cultures. Subsequently, amino acid production and metabolomic activity of glycolytic intermediates are increased when compared with monolayer substrates of the same cell line [22],[23].

A wide array of scaffolds can be used to recapitulate the TME and support differentiation of 3D culture, given that TME is pivotal for the regulation of a diverse array of processes including, migration, proliferation, differentiation, and cell-cell communication [24]. Often interchangeable within the literature, spheroids and organoids differ in complexity. Typically, spheroids are rounded and are comprised of cells grown initially in 2D, and as such retain some simplicity of gene expression. Growth is often achieved using hanging drop method or an ultra-low attachment plate and is ideal for the study of diffusion gradients and core hypoxia [25].

Given the current trajectory of 3D cancer models and their appeal to support the reduction of animal research, it is therefore safe to assume that a complex OvCa on a chip model will soon be achievable. This meta-analysis aims to evaluate the current landscape of OvCa cell models to elucidate differences presented in their genetic profile and associated signalling pathways, when grown in 3D compared to 2D monolayer culture.

## 2. Materials and Methods

### Study Design

The review was designed with the intent to search current literature for studies modelling OvCa using 3D culture techniques and assess the differences in gene regulation between 2D and 3D cultures. The National Centre for Biotechnology Information (NCBI) PubMed data base was searched for studies relevant to the scope of the review between the years 2012 and 2022. No limitations to original language were applied, as long as English translations were available. The filter for human studies was utilised. Search terms applied include: “cancer” AND “ovar*” AND “3d” NOT “sound” NOT “ultra” NOT “imaging” NOT “Ultrasound” NOT “Review”. Literature that was inaccessible via the university institutional access were also removed. Additional searches through NCBI, Sequence Read Archive (SRA) and Gene Expression Omnibus (GEO) accession platforms were also utilised.

Inclusion criteria: Studies were included if they encompassed 3D OvCa models as well as 2D comparisons. In addition, those with associated data from sequencing arrays and RNA sequencing, accessible through GEO or SRA, were also sought.

Exclusion criteria: Studies were discarded if they did not meet the original search criteria. Additional studies that were excluded comprised of those with a lack of comparative 2D culture, no open access and no human samples i.e., the use of animal (usually murine) cell lines. Final exclusion criteria for enrichment encompassed studies with no associated data.

### Cell Culture and 3D modelling

Unless otherwise stated all reagents were purchased from Thermofisher Scientific. The serous ovarian adenocarcinoma cancer cell line SKOV-3 (ECACC 91091004) were seeded in conventional culture-treated polystyrene T75 flasks. Cells were grown in Dulbecco modified eagle’s medium (DMEM), supplemented with 10% foetal bovine serum and 1% penicillin-streptomycin. Media changed every 2 – 3 days with experimental work proceeding after 3 passages. Cell suspension concentrations were calculated using trypan blue exclusion method. For monolayer substrate comparison, cells were seeded in triplicate, at a density of 5×10^6^ in an Ibidi 8-well chamber (Ibidi, Munich, Germany) with complete medium. 3D cultures were generated using a 1:12 ratio of cells suspended in medium mixed with GelTrex™ (batch: 2158356). Each well contained a final concentration of 300μl. The chamber was left to incubate at 37°C for 30 minutes to allow for gelation, 100μl of media was then added to each well. Media changes took place every 2 – 3 days up to day 10. Images were captured each day using a Nikon TS100 Inverted Phase Contrast light microscope (Nikon, Tokyo, Japan).

Certificate of analysis and declaration of mycoplasma free cultures were provided upon receipt of cells from PHE and validated in house with DAPI staining; cells were used following 3 passages from purchase.

### Immunofluorescent imaging

On day 10, media was removed. Both 2D and 3D cultures were fixed with 4% paraformaldehyde in PBS for 10 and 30 minutes respectively. Chambers were washed x3 with PBS following incubation with 0.1% triton-x, for 10 minutes. Chambers were again washed prior to blocking with 10% bovine serum albumin (BSA) (Sigma Aldrich, Burlington, MA, USA), for 1 hour at room temperature. BSA was then removed for phalloidin (ATTO-TEC, Siegen, Germany) actin staining, using a 1:1000 dilution in 1% BSA for 30 minutes at room temperature. Chambers were again washed x3 with PBS before the administration of a final DAPI (Invitrogen, Massachusetts, USA) nuclear stain for 10 minutes. Samples were washed to remove residual DAPI and kept hydrated in PBS prior to imaging.

### Laser Scanning Confocal Microscopy

Laser scanning confocal microscopy (LSM780, Carl Zeiss, Oberkochen, Germany) was used for 3D imaging of cells cultured in a glass substrate and encapsulated in 3D Geltrex hydrogel. The cell samples were subject to excitation\emission wavelength at 405 nm\410 nm-495 nm and 488 nm\495 nm – 620 nm, for imaging of nuclei (DAPI) and actin (phalloidin), respectively. The emitted fluorescence signal was recorded using photomultiplier tube (PMT) detectors. The optical Z-stacks were acquired using 63x objective (A plan-Apochromat 63x/1.4 Oil immersion, Carl Zeiss). The laser power, detector gain, and scan speed were optimized to avoid photobleaching. The image size was 2048 pixels x 2048 pixels, with a voxel size of 40 nm x 40 nm in the XY-plane, and 250 nm in the Z-direction. The images were deconvoluted using automatic deconvolution mode with theoretical point spread function using Huygens Essential software (Scientific Volume Imaging, The Netherlands). Avizo software (Thermo Fisher Scientific, Waltham, MA, USA) was used for 3D visualization.

### RNA Sequencing – Sequence Read Archive (SRA)

NIH Sequence Read Archive (SRA) data were found using the same search terms outlined in the study design. SRA data in the form of RNA sequencing reads produced with Illumina NextSeq 500 and Illumina HiSeq 2500 were acquired for re-analysis, accession IDs are outlined below in table 2. Briefly, relevant data in the form of FASTQ files were transferred from the SRA data base via Amazon Web Services for in house analysis (Table 2) – full list can be seen in (Supplementary Table 1). The corresponding scaffold used within each study are as follows. PRJNA472611, 3D cells were embedded within agarose; PRJNA564843 cells were grown upon a layer of onmental fibroblasts embedded within Collagen; PRJNA530150 3D cells were grown in Matrigel.

**Table 2.**
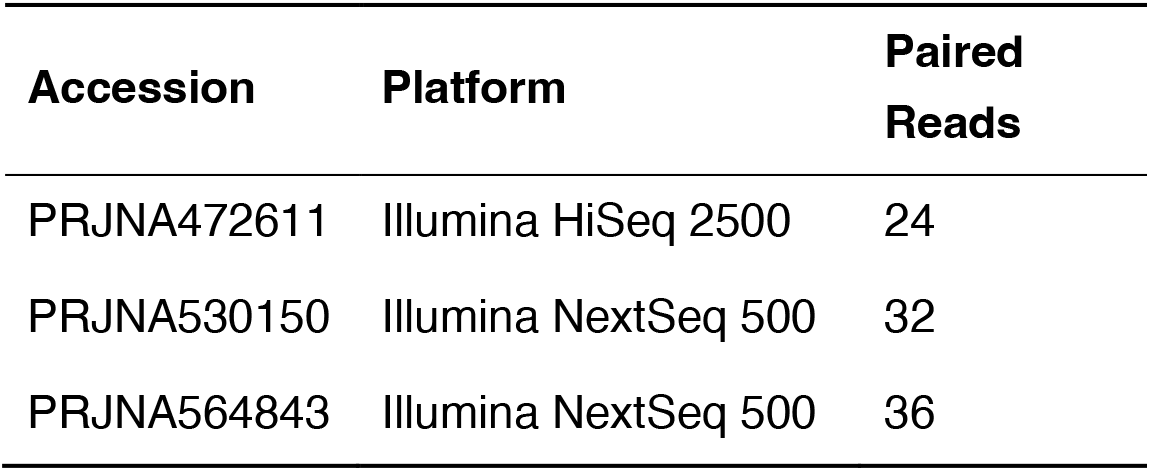
Accession codes from RNA sequencing of 2D and 3D OvCa cell models.

The raw RNAseq data was produced using the pipeline previously described to standardise the results for comparison [26]. Briefly, TopHat2 (v.2.1.1) was applied to align reads to the reference human genome, GRCH38 (hg19) using the ultra-high-throughput short read aligner Bowtie2 (v.2.2.6). Where applicable replicates were merged according to a selection criterion taking only high-quality mapped reads (<30), using Samtools (v.0.1.19). Subsequent transcript assembly and quantification followed using Cufflinks (v.2.2.1). Finally, differential expression profiles were obtained for further analysis using Cuffdiff (v.2.2.1).

### RNA Sequencing – Statistical Analysis

The expression data was analysed in R (v. 4.1.0, The R Foundation for statistical Computing, Vienna, Austria) with R studio desktop application (v.2022.07.2, RStudio, Boston, MA, USA) using specific libraries for modelling, visualisation, and statistical analyses for the identification of differentially expressed genes (DEGs). Similar to our previous work, Pearson correlation coefficient was applied for the estimation of gene expression patterns and student’s t-test was utilised to assess statistical significance between expression profiles (i.e., 2D vs 3D). Significance thresholds were set for a p-value < 0.05. For identification of enriched pathways in omics data pathfindR was employed. Volcano plots for visualisation were generated using R package ggplot2 (v.3.3.5). DEGs were identified and isolated for subsequent enrichment analysis. Furthermore, we have used the OmicsPlayground online application for exploring the transcriptional landscape of ovarian cancer cells grown in 2D and various 3D systems using as scaffolds agarose, collagen and Matrigel [27].

### Gene Expression Omnibus (GEO) Array – Statistical Analysis

Genomic data sets (accession numbers: PRJNA232817 and PRJNA318768) were downloaded from NCBI public repository GEO archive. These OvCa cells were grown using ultra-low attachment and hanging drop techniques. The GEO2R web application was accessed to re-analyse the expression data in line with the research questions within this study (control 2D samples vs. control 3D samples). Thresholds were again set at p-value < 0.05 and LogFC2 > 1 with applied Benjamini & Hochberg (False discovery rate). Volcano plots were generated through GEO2R (https://www.ncbi.nlm.nih.gov/geo/geo2r/).

### Functional Enrichment Analysis

Differentially expressed genes (DEGs), identified through GEO2R and SRA analysis, were then subjected to functional enrichment analysis. Funrich (v.3.1.3), was accessed to provide a functional annotation including associated sites of expression, biological processes, and pathways. Enrichment Analysis was performed using Omics Playground for the functional comparison of OvCa genes in 2D vs. 3D [27].

### Presentation of Data and Statistical Analysis

Global distribution infographics were generated using R (v.4.1.0) with R studio (v.2022.07.2) along with ggplot2 (v.3.3.6), maps (v.3.4.0) and world map data from natural earth (0.1.0). Subsequent comprehensive background analysis and graphs pertaining to publication data, cell line frequency and associated characteristics were generated using GraphPad Prism9^®^ (v.9.4.1 - GraphPad Software, Inc.). Statistical reliability of Omics Playground data are ensured through the incorporation of Spearman rank correlation, GSVA, ssGSEA, GSEA and Fisher extract test [27].

## 3. Results

### 3D Ovarian Cancer models

#### Literature overview

The geographical spread of the fifty studies selected suggests that the United States of America (USA) are the top publishers of 3D OvCa modelling with over 50% of the research accessed originating within the USA. China, Italy, Korea and the UK follow, with the majority of the work originating from Europe or North America (Figure 2A, B).

**Figure 1.**
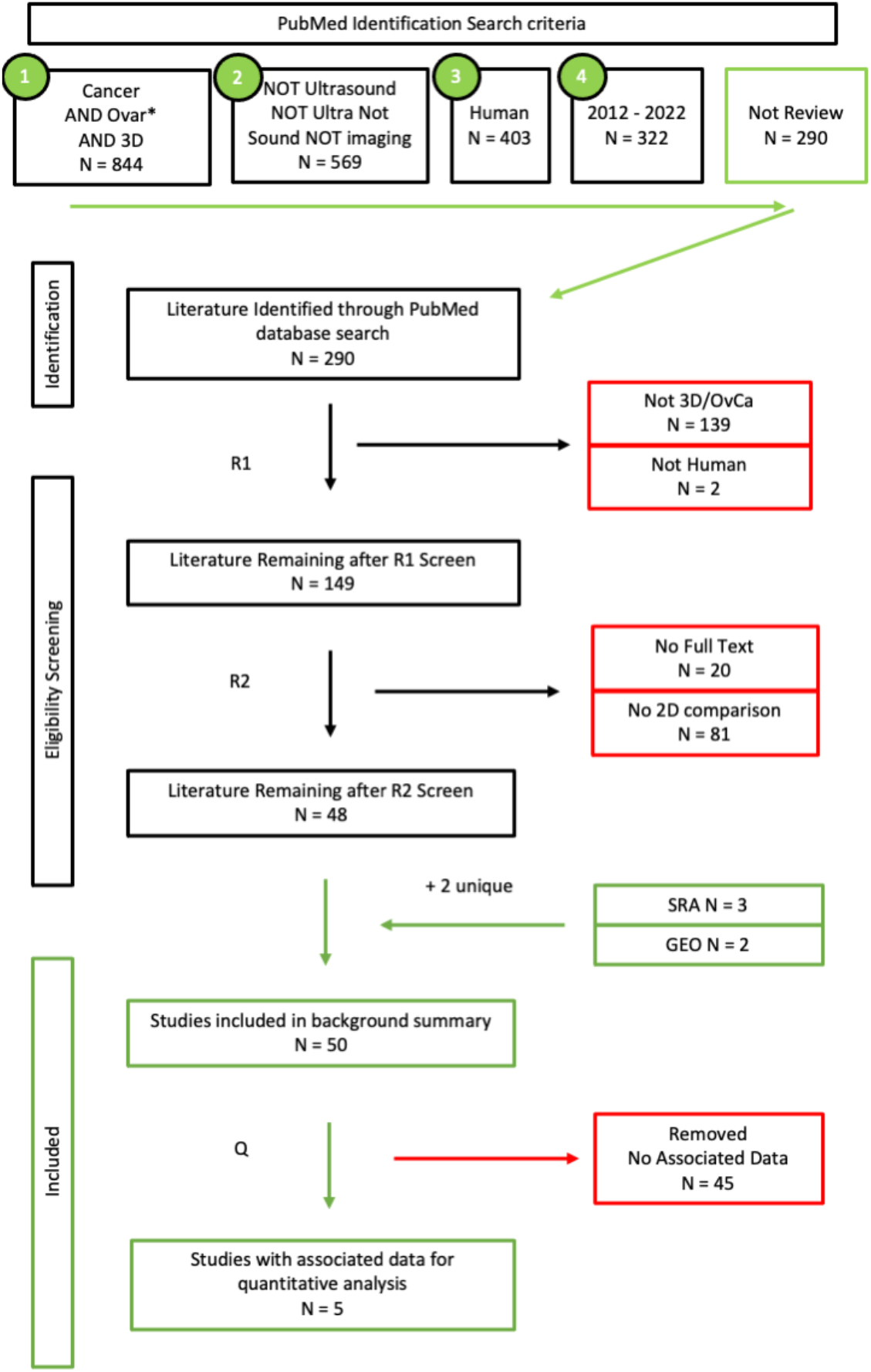
Search Criteria workflow. Studies accessed through Pubmed.gov on the 25/06/2022. Using predefined search terms. Articles were subjected to 2 rounds of screening by two independent reviewers. Additional data sought through Sequence Read Archive and Gene Expression Omnibus 07/2022. Studies were split into two groups: those suitable for the background summary (N = 50) and those containing associated data (N = 5).

**Figure 2.**
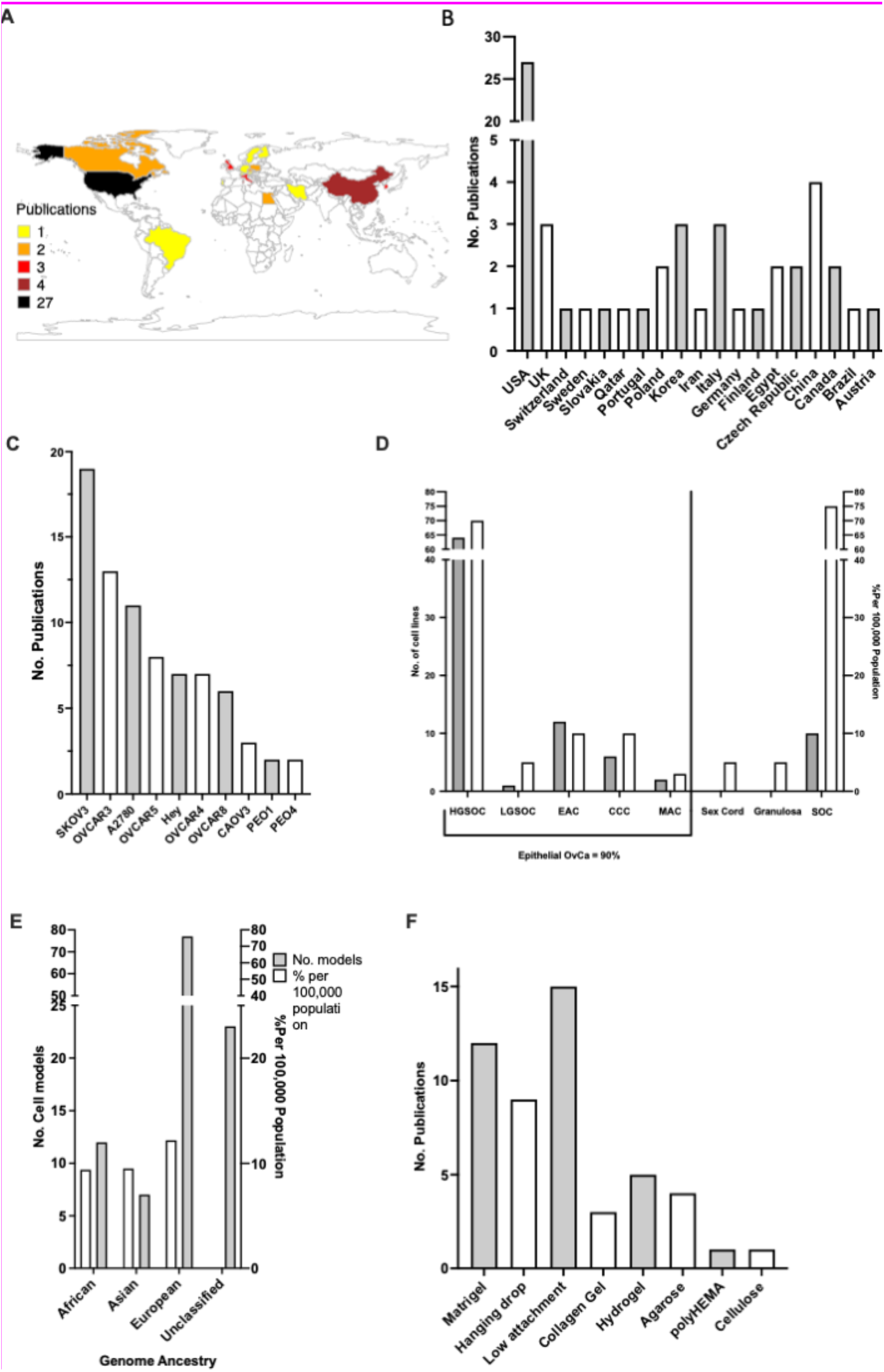
**(A)** Gradient map depicting the global spread of publications 2012 – 2022; **(B)** Chart showing no. of publications per country 2012 – 2022; **(C)** Top cell lines used in 3D Ovarian cancer (OvCa) within the literature; **(D),** Trends between the distribution of cell models against actual global rates (white) pertaining to OvCa subtype (grey); **(E),** Genome ancestry of cell lines used (grey), contrasted with actual global OvCa ethnicity rates (white) (2012 – 2022); **(F)** The ten most frequently used scaffolds for supporting growth of OvCa cells (circa 2012 - 2022) selected from the publication corpus analysed.

To achieve 3D culture, cell lines are grown within a fabricated ECM also known as a scaffold. Within the literature the most commonly used scaffolds for 3D OvCa growth were pre-coated low attachment plates, followed by Matrigel, hanging drop method and plant-based hydrogel (Figure 2C). Over 43 unique OvCa cell lines were utilised throughout the studies. The top 10 represent an array of OvCa subtypes (Figure 2D). The ovarian carcinoma cell line SKOV-3 was the most frequented within the literature, appearing on 19 instances. The trend of studies focusing on OvCa subtypes was compared with the actual global incidence rates. For epithelial OvCa the cell models used followed a similar trend in frequency to actual global incidences, with HGSOC being the most prevalent form of OvCa and also the most studied. Of note sex cord stromal and granulosa OvCa comprises 10% of global cases, however no 3D models were found within the studied literature. The genome ancestry of the cell lines is often overlooked, however given the disparity in care the background of the cell lines used was also sourced (Figure 2E). A disproportionate number of cell lines used are either White (N = 80) in origin or are considered unclassified i.e. no available data (N = 30).

### Differentially Expressed Genes

Data accessed through SRA and GEO were screened for OvCa cells grown in 2D and 3D under similar conditions. Three separate studies were chosen encompassing 19 cellular models grown under normal conditions in agarose, Matrigel and collagen-based scaffolds. All cell lines grown in 3D showed differential gene expression when contrasted with the same cell lines under the same conditions but grown in 2D (Figure 3). The number of statistically significant differentially expressed genes (DEGs) with p < 0.05, between the 2D and 3D cultures ranged between 234 in PEO1, to 1429 in OVCAR5 cell line.

**Figure 3.**
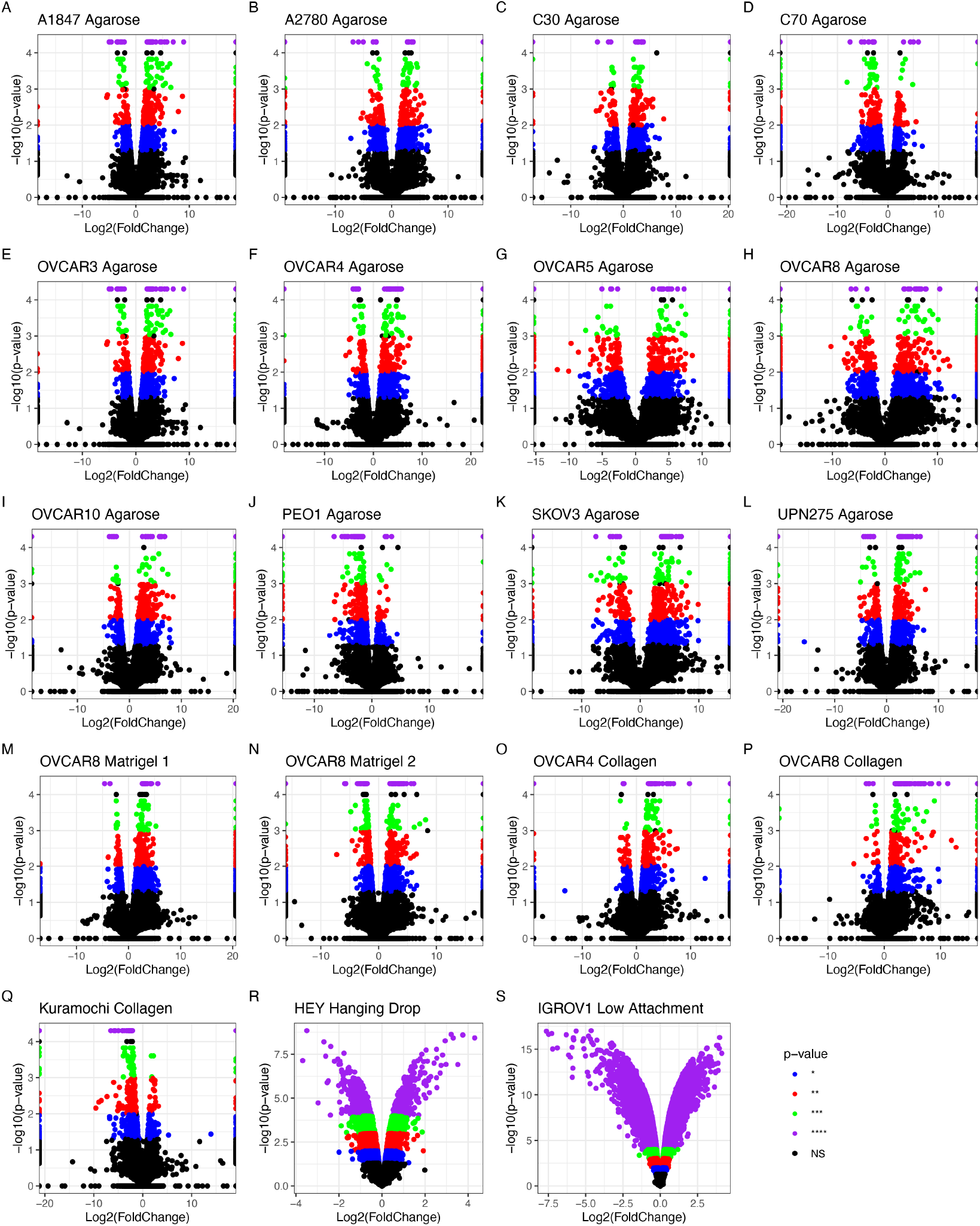
Differentially expressed genes (DEGs) detected by RNA sequencing analysis of OvCa cell lines grown in 2D contrasted with 3D. **(A)**–**(Q)** show data extracted from RNAseq experiments **(R)**–**(S)** show data extracted from microarrays. Significance thresholds for **(A)**–**(Q)** are set at NS > 0.05 = grey/black, *p < 0.05 = blue, **p < 0.01 = red, ***p < 0.001 = green and ****p < 0.0001 = purple. **(R)**–**(S)** p-value threshold = 0.05, NS data is shown in black. **(A)** – **(L)** have agarose as scaffold, **(M)** – **(N)** are Matrigel, **(O)** – **(Q)** are collagen, **(R)** is hanging drop and **(S)** is low attachment. **(A)** A1847 - Endometrioid Carcinoma of the Ovary (EAC); **(B)** A2780 - EAC; **(C)** C30 - carcinoma; **(D)** C70 - carcinoma; **(E)** OVCAR3 - HGSOC; **(F)** OVCAR4 – HGSOC; **(G)** OVCAR5 - HGSOC; **(H)** OVCAR8 - HGSOC; **(I)** OVCAR10 - HGSOC; **(J)** PEO1 - HGSOC; **(K)** SKOV-3 - Carcinoma; **(L)** UPN275 - Mucinous adenocarcinoma (MAC); **(M)** Kuramochi - HGSOC; **(N)** OVCAR4 Collagen - HGSOC; **(O)** OVCAR8 Matrigel 1 - HGSOC; **(P)** OVCAR8 Matrigel 2 - HGSOC; **(Q)** OVCAR8 Collagen - HGSOC. **(R)** HEY – HGSOC; **(S)** IGROV1 – EAC.

The HGSOC OVCAR8 appeared in all three studies with different accompanying scaffolds: Matrigel, agarose and collagen. Therefore, additional analysis explored the effects of different scaffolds on the genetic profile of these cells (Figure 4). All conditions influenced differential regulation of OVCAR8’s transcriptional profile. 13 DEGs were identified (Table 3) based on their common dysregulation between scaffolds when grown in 3D. Similarly, these genes were seen to feature highly throughout the other 3D models i.e., dysregulation of ANGPTL4 appeared in 12/19 of the studies. When comparing DEGs identified between OVCAR8 cells grown in 2D and 3D, eight were found to be common regardless of their scaffold type (Figure 4 and Table 3).

**Figure 4.**
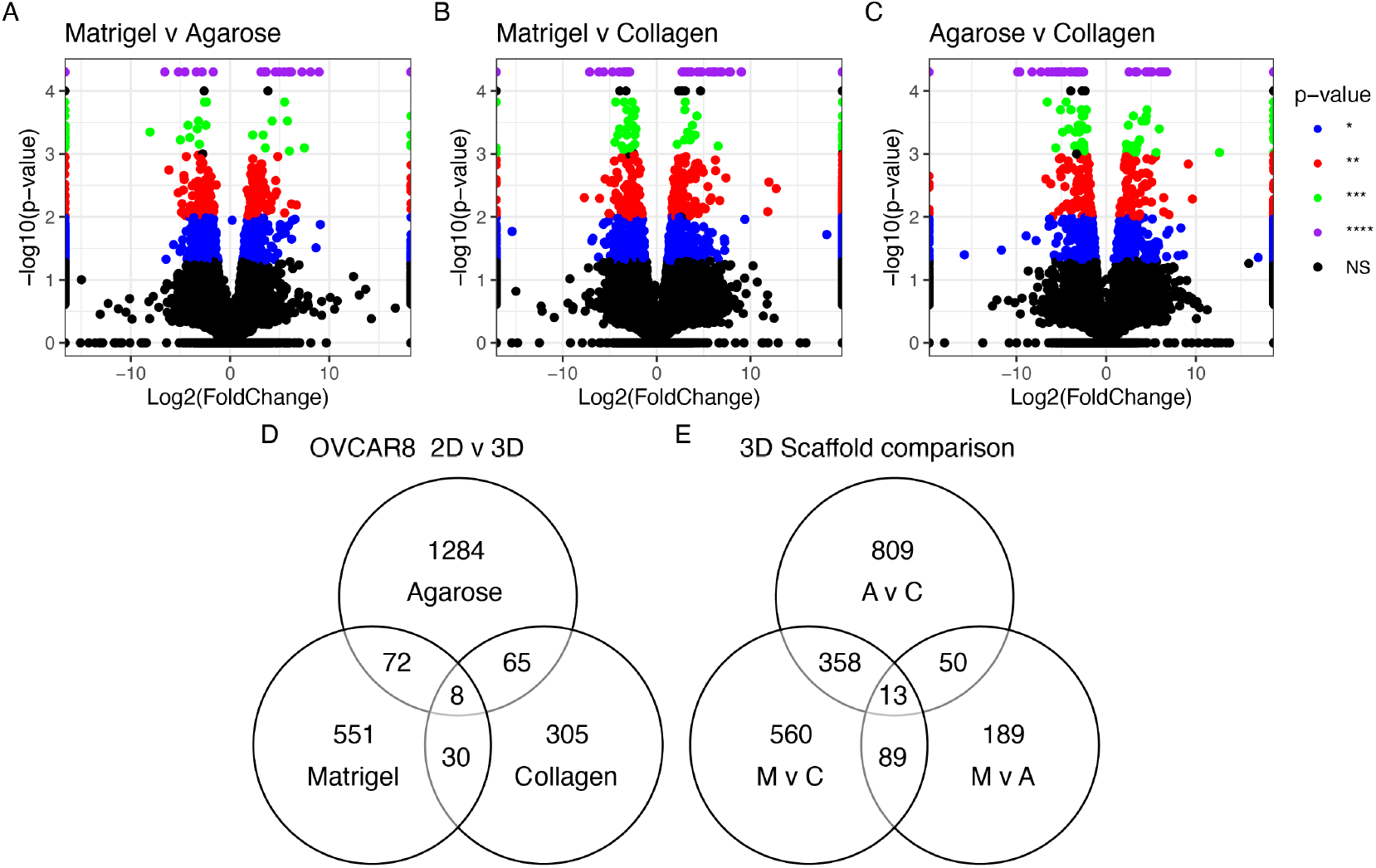
Differentially expressed genes seen in OVCAR8 grown in 3D**. (A)** Agarose vs. Collagen; **(B)** Matrigel vs. Agarose; **(C)** Matrigel vs. Collagen. Threshold set at p < 0.05. **(D)** Common genes between **(A - C); E)** Common genes seen between OVCAR8 grown in 3D vs. 2D. M: Matrigel, C: Collagen, A: Agarose.

**Table 3.**
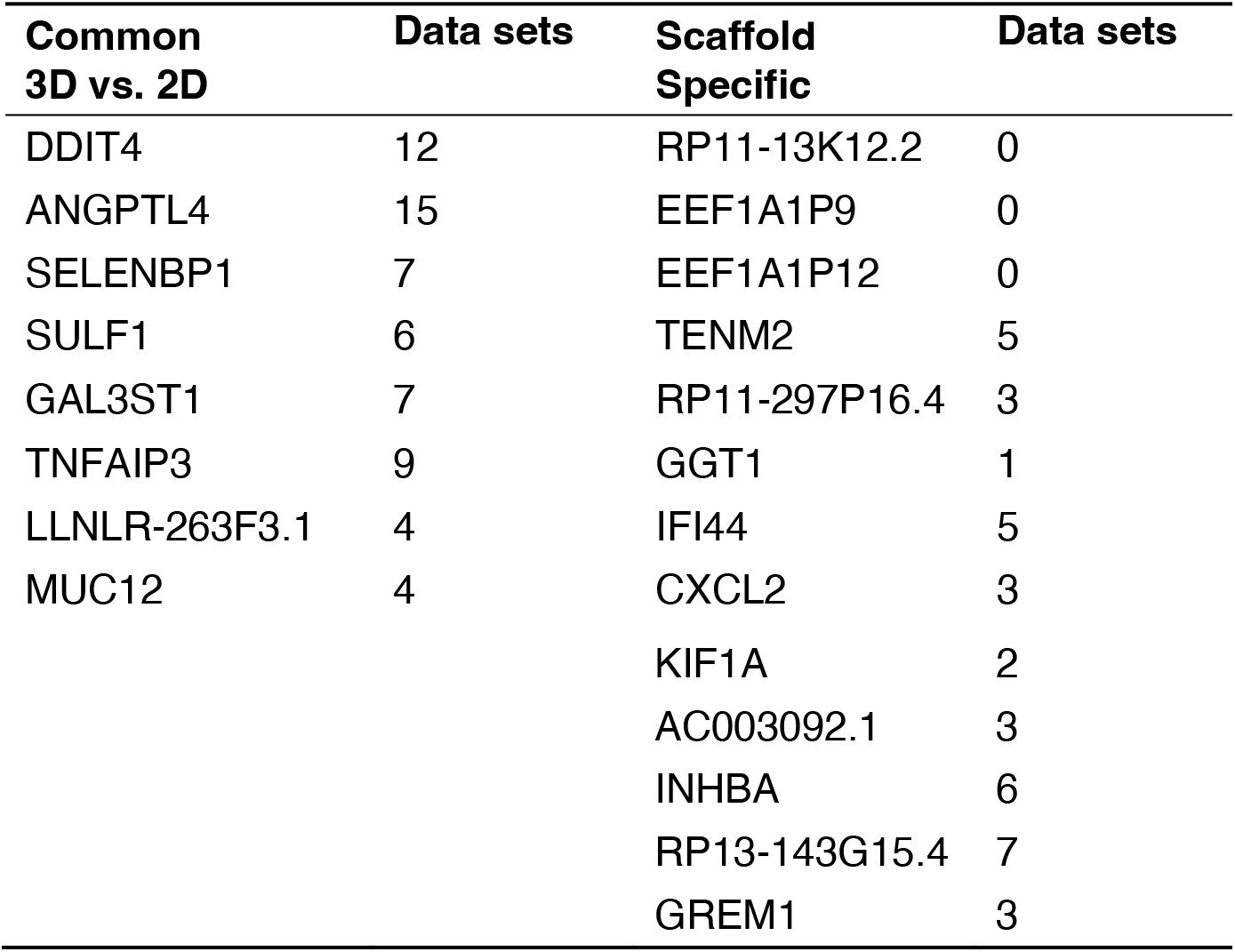
OVCAR8 genes commonly differentially regulated in 3D conditions grown on agarose, collagen and Matrigel compared to 2D cultures

### The impact of scaffold and 3D set up as compared to 2D culture on the genetic profile of OvCa cells

We explored the transcriptional landscape in 2D and 3D cultures in 3 different scaffolds (agarose, collagen and Matrigel) for the OVCAR8 cell lines.

The cells grown in 3D on Matrigel, agarose and those grown on a basement layer of normal omental fibroblasts embedded within collagen, were compared with standard 2D monolayer cultures (Figure 5). The expression profiles of the top 150 DEGs with respect to growth conditions are shown in Figure 5A (Supplementary Table 2). This gene set shows a large variability across the four growth conditions. Initial observations reveal a high degree of similarity in gene expression between samples grown in agarose and Matrigel. Collagen samples however show an expression profile that diverges from the 2D expression profile to a lesser extent than OVCAR8 grown on other scaffolds. T-SNE analysis (Figure 5B) recapitulates these observations showing a partial clustering of the 3D profiles, with the collagen 3D culture standing out and showing the highest level of similarity with the 2D culture experiments. The top functional groups of the differentially regulated genes included key metabolic pathways such as glycolysis (Figure 5C).

**Figure 5.**
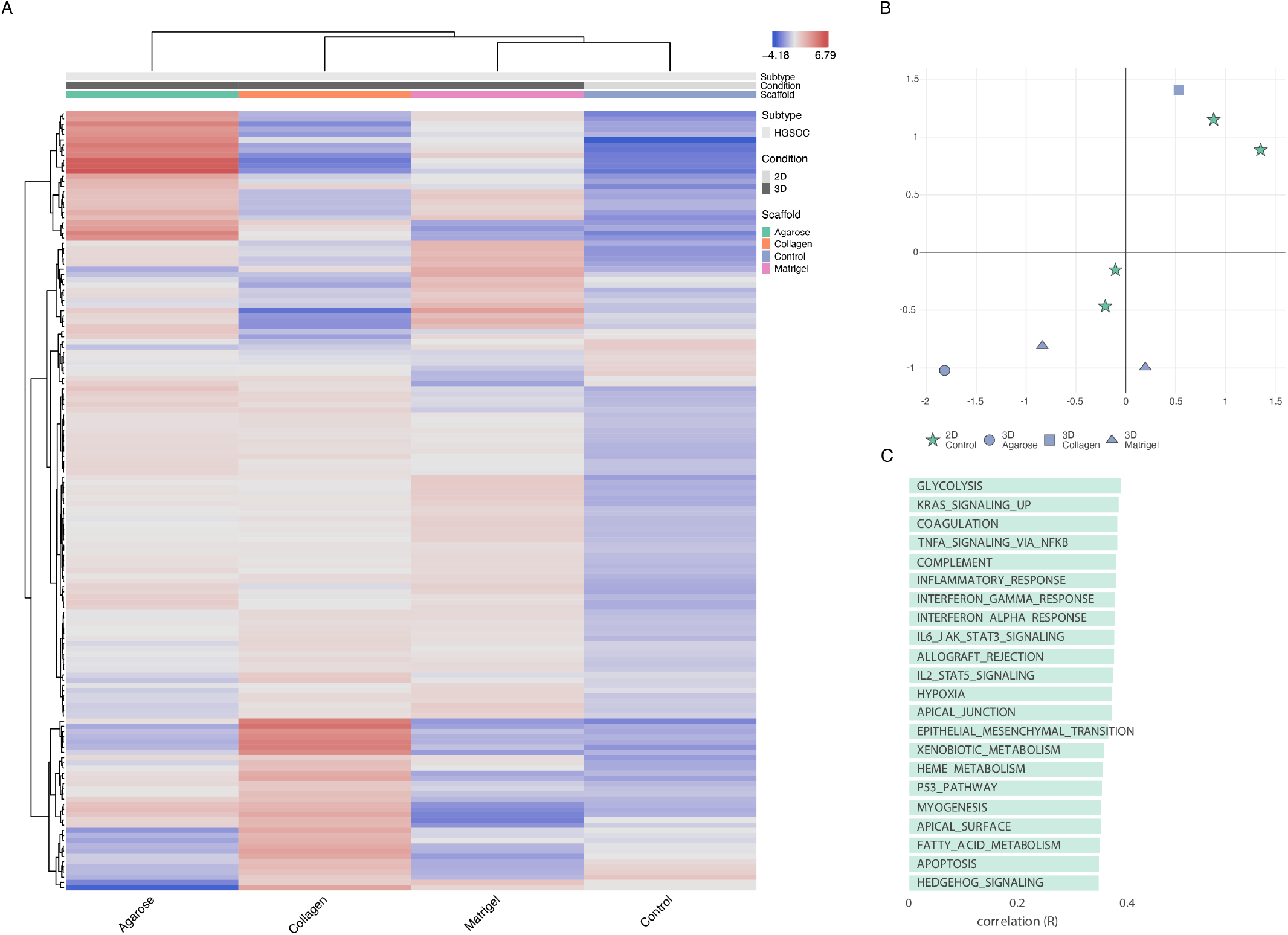
OVCAR8 transcriptional profile in 2D v 3D. **(A)** Top 150 differentially regulated genes from OVCAR8 grown under 2D and 3D conditions. Data originating from 3 unique studies, encompassing 4 growth conditions. 3D cells grown in Matrigel, Collagen and Agarose. 2D cells grown under standard lab conditions as matched controls to each 3D experiment. The gene name list is available in Supplementary Table 2; **(B)** T-SNE plot of the genetic profiles of the HGSOC OVCAR8 grown in Matrigel (at 7 and 14 days - triangle), Collagen (square), Agarose (circle) and Monolayer (stars); **(C)** Functional analysis of the top 150 differentially regulated genes between 2D and 3D growth conditions showing key biological pathways associated with them.

Next, we explored the genes transcriptional signatures in the three scaffolds and in the 2D control experiments. We clustered the genes based on pairwise coexpression scores and visualised them using a uniform manifold approximation and projection dimensionality reduction technique (UMAP) (see Figure 6A). We found localised phenotypic clustering patterns in OvCa embedded in collagen and agarose with less variance in phenotypic expression recorded for samples grown in Matrigel, when compared with 2D. Moreover, Matrigel culture showed an inverted gene expression signature compared to 2D control experiments. Similarly, we analysed cancer hallmark sets with the DEGs of OVCAR8 grown in 2D compared to 3D data (see Figure 6B). Processes with high covariance include: K-Ras signalling, angiogenesis, interferon alpha and gamma response, TNF alpha signalling as well as epithelial to mesenchymal signalling.

**Figure 6.**
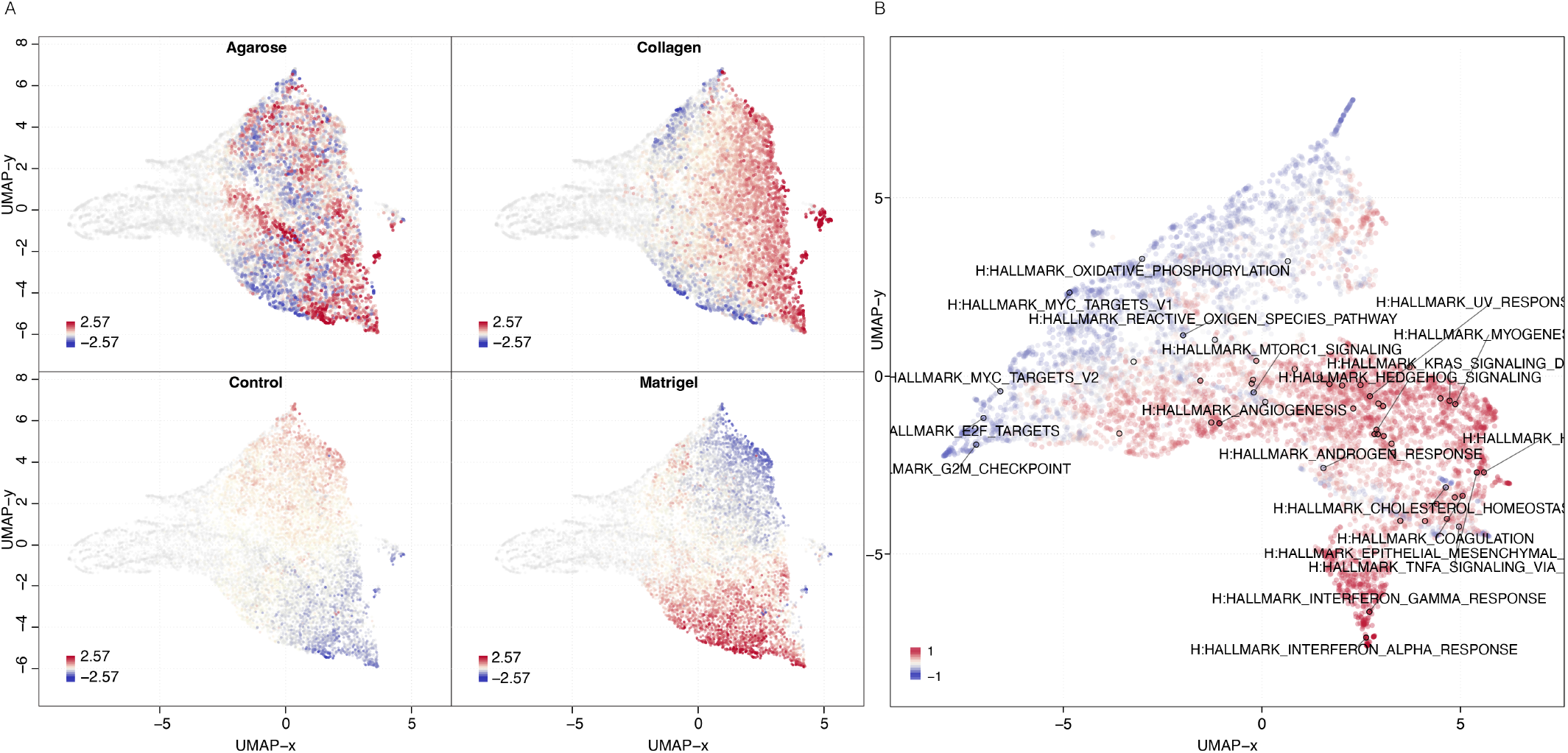
Gene and phenotypic hallmark signature profiles. **(A)** UMAP clustering of genes coloured by relative log-expression in four growth conditions: agarose, collagen, Matrigel and 2D controls. The distance metric is covariance. Genes that are clustered nearby have high covariance. **(B)** UMAP hallmark covariance using OVCAR8 grown in 2D and combined 3D data. Clustering of associated hallmarks. Processes upregulated in 3D are indicated in red. Downregulated are indicated in blue.

### Functional Enrichment – 2D vs. 3D

A panel of genes were identified as commonly disregulated in 3D cultures compared to 2D growth conditions. The cumulative 3D data encompasses OVCAR8 grown on Matrigel, agarose and collagen, while the control data is composed of the experiments using 2D growth conditions. The following genes showed statistically significant differential expression (p < 0.05): C3, CXCL1, CXCL8, IL1B, SLPI, FN1, IL6, DDIT4, PI3, LAMC2, CCL20, MMP1, IFI27, CFB, ANGPTL4 and CXCL2 (Figure 7).

**Figure 7.**
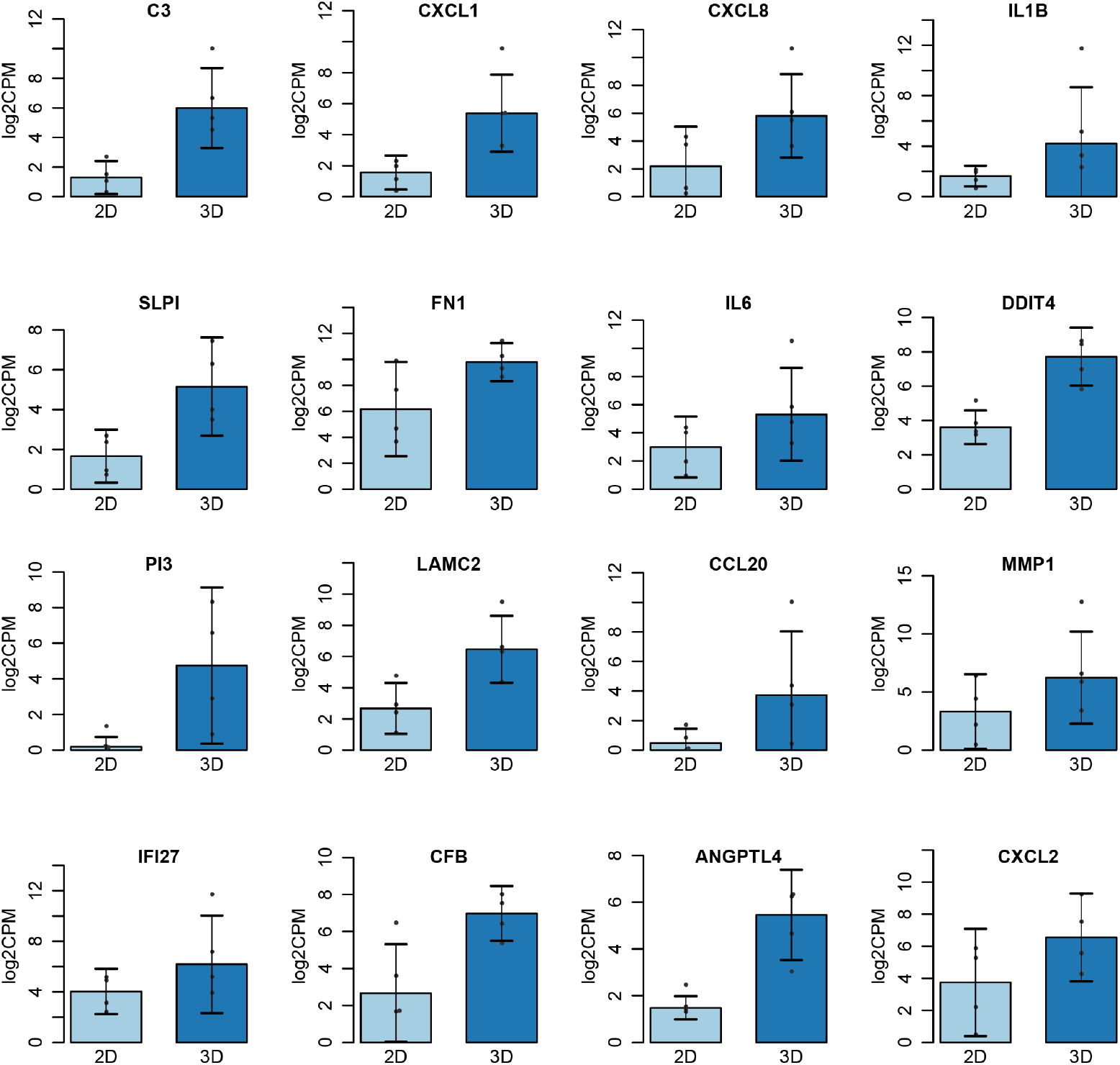
Top Genes Differentially Expressed 2D vs. 3D. Cumulative data for 3D taken from OVCAR8 embedded within Matrigel, Agarose and Collagen. Significance threshold *P < 0.05.

### Scaffold Specific Biomarkers – 2D vs. 3D

Next, we examined the transcriptional landscape to identify potential biomarkers of growth conditions (Figure 8). For this we have used a variety of machine learning algorithms as implemented in the OmicsPlayground v2.8.10 to compute a cumulative importance score for all DEGs. The results highlighted 8 key genes that can be used as predictive scaffold biomarkers (Figure 8A). Specifically, cells grown in agarose show condition specific expression for 4 genes: C3, MMP1, IL1B and CCL20. Three potential markers of cells grown in collagen were identified namely: the interferons IFI44L and IFI27 as well as COL3A1. Matrigel was represented with only one significant growth marker: DDIT4. While these 8 biomarker candidates show the highest importance scores, a variety of other genes show scaffold specific expression as well (Figure 8H), suggesting that a number of gene panels can be created to evaluate the impact of growth conditions on the genome biology.

**Figure 8.**
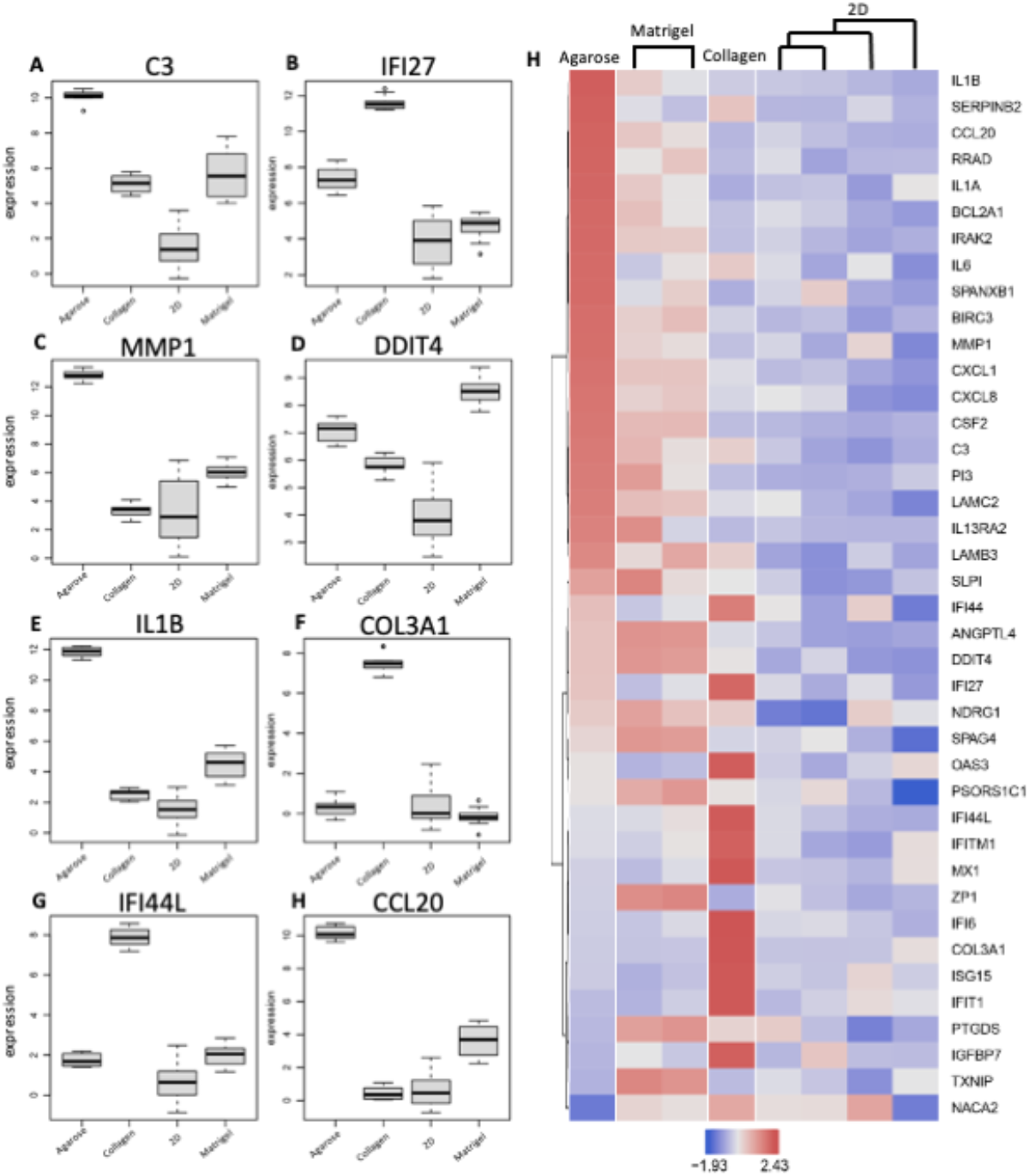
Scaffold specific biomarker identification. **(A - H),** The top 8 genes implicated with expression specific profiling for each condition; **(I)**, Biomarker Heatmap: expression heatmap of top gene features according to their variable importance score. Importance scores are calculated based on multiple machine learning algorithms including LASSO, elastic nets, random forests, and extreme gradient boosting.

### Cell line specificity impact on scaffold selection

Following the analysis of the impact of scaffold and the 3D v 2D environment on the transcriptional landscape of the ovarian cancer cell line we looked at differential expression patterns between various cells lines grown on agarose and collagen scaffolds. As expected, we found a good separation of the cell line gene expression characteristics on both scaffolds (Figure 9A,B) using the top 150 differentially expressed genes. Most cell lines have also shown a fair discrimination between the 2D and 3D cultures on agarose, and also a good segregation between cancer subtypes (Figure 9C). However, A1847, OVCAR3, OVCAR4 and SKOV3, on agarose and all cells on collagen (Kuramochi, OVCAR4, and OVCAR8) show poor differentiation between the growth conditions suggesting that these scaffolds are potentially not optimal for recapitulating the tumour environment more accurately than classical 2D cultures in these cell lines.

**Figure 9.**
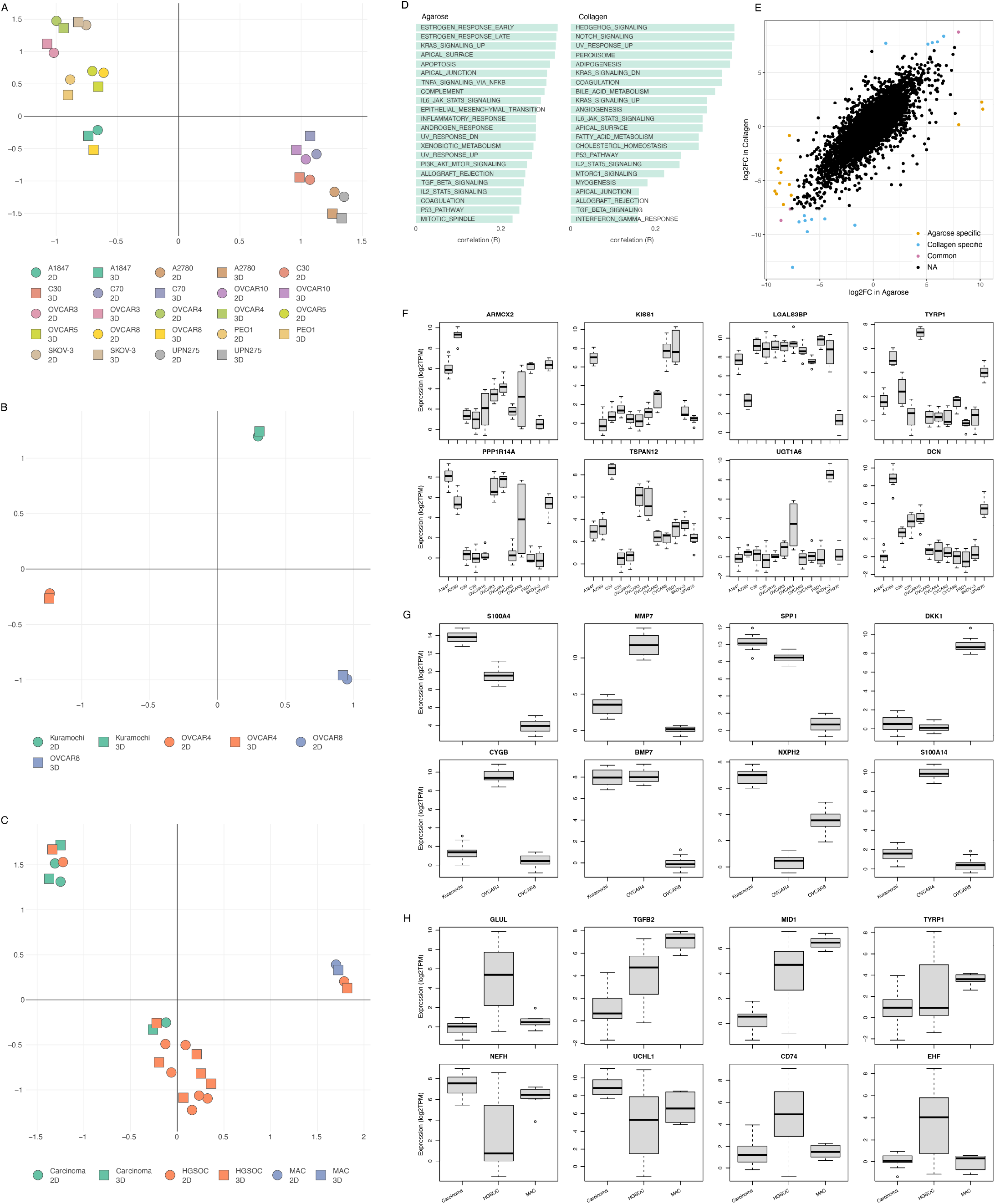
Cell line specific transcription in agarose and collagen. **(A)** and **(B)**, T-SNE plot of the genetic profiles of cell lines grown in agarose and collagen respectively against a 2D control. **(C)** Umap plot of the transcriptional profile of cancer subtypes in agarose vs 2D control, **(D)** Functional analysis of the top 150 differentially regulated genes between 2D and 3D growth conditions showing key biological pathways associated with them for agarose and collagen, **(E)** Similarity of gene differential expression in OVCAR4 v OVCAR8 in collagen versus agarose, **(F) – (H)** The top 8 environment biomarkers for cell lines grown in agarose **(F)** and **(H)**, and collagen **(G)**.

Functional analysis reflects the diversity of the cell lines grown on each scaffold (Figure 9D). With sex hormones specific pathways characterizing the agarose cultures while cell growth and development pathways, as well as fatty acids metabolism being the dominant features of the collagen grown cell lines. The scaffold impact on cell line specificity was explored by comparing the differentially expressed genes between OVCAR4 and OVCAR8 in agarose and collagen (Figure 9E). We found that there is a good level of correlation between gene expression fold change in the two cell lines for agarose and collagen. Of the top differentially expressed genes, three, SLC34A2, LY6K, BMP7, show the same level of dysregulation between OVCAR8 and OVCAR4 in both growth conditions. However, we also identified 13 genes that show a scaffold specific differential expression pattern between the two cell lines: MMP7, LAMA3, IGFL1, S100A14, ELF3, CYGB, ITGB6, DKK1, TACSTD2, IL7R, LGALS13, IFI6, FOXD1 being collagen specific, and IL1B, MMP1, CP, UBB, NUPR1, SCGB2A1, GPNMB, IGFBP2, GDF15, CCL20, CYP1A1, VTCN1, KRT19 agarose specific.

Finally, the differential expression patterns identified a number of genes that show both a cell, tumour subtype, and scaffold specific behaviour and can be used as environment biomarkers (Figure 9F-H).

### Recapitulation of 3D OvCa using GelTrex

Leveraging the lessons learned from the study of the transcriptional landscape of OvCa cell lines in different conditions, we attempted to capture the changes in the phenotype between the 2D and 3D cultures. For this we grown SKOV-3 cells, in 3D using the hydrogel-based scaffold GelTrex^™^. Hydrogel was chosen as it encompasses one of the most common scaffolds within the literature and is not animal derived. In addition, this work sought to assess the ease of using non-established methodology for in house recapitulation. As such hanging drop and ultra-low attachment plates were not included as their use with OvCa is well established within the literature.

Figure 10 shows the growth of cells over the course of a 9-day period. Here we adopted a simplistic approach and used a previously tried and tested gel known as GelTrex. Following the embedding process cells began to aggregate and form spheroid like structures [28]. These structures-maintained circularity and continued to expand in volume as time progressed. The results suggest that the changes at genomic level have a direct impact on the 3D aggregation of cells.

**Figure 10.**
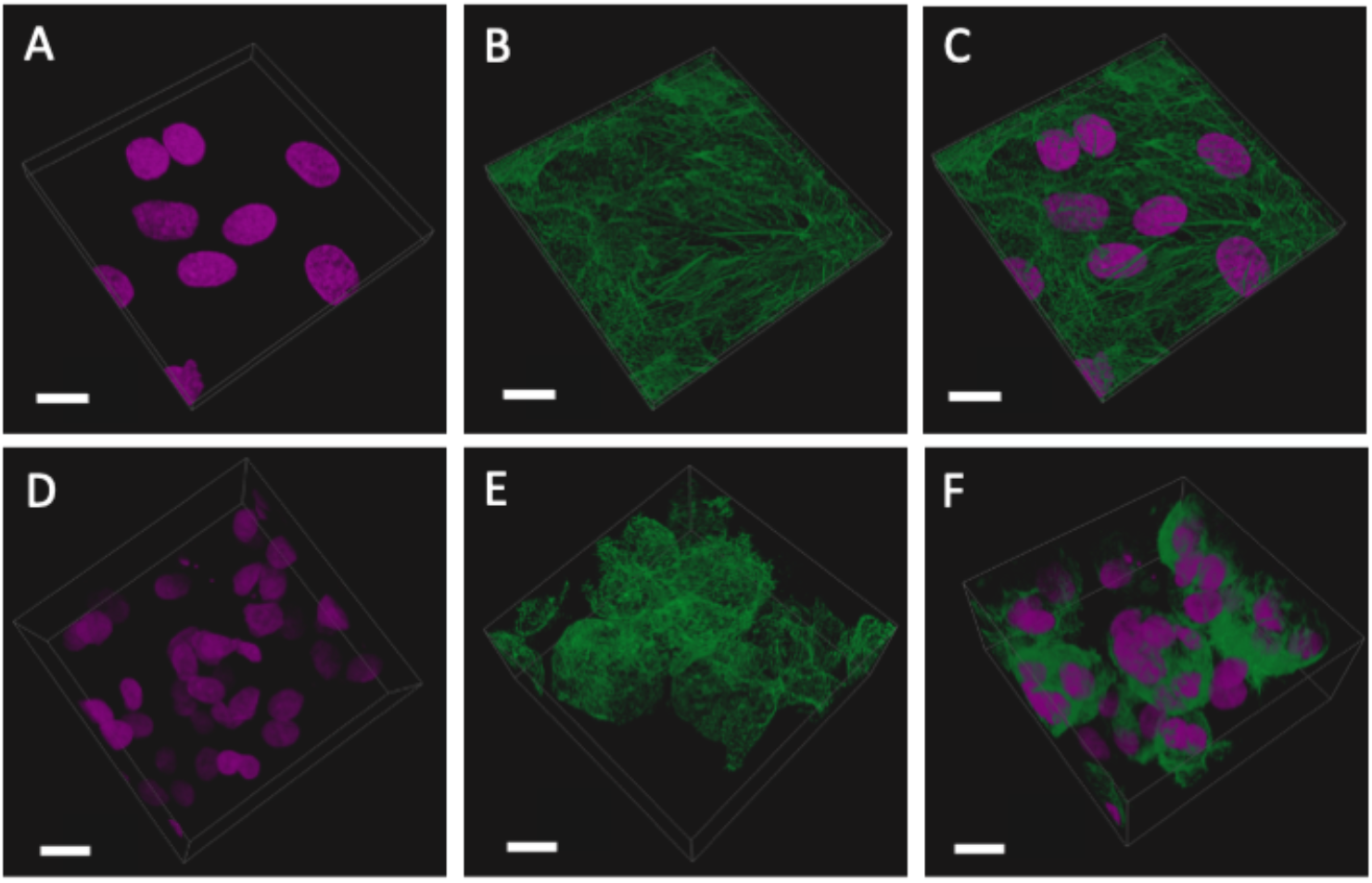
SKOV-3 cells grown for 9 days in conventional monolayer formation compared with those em-bedded in GelTrex^™^. **(A – C)** Monolayer cells: nuclei (pink), phalloidin (green) and overlay, showing a single plane of cells across a flat glass substrate; **(D – F)** 3D cells: nuclei (pink), phalloidin (green) and overlay, showing aggregated spheroids with multiple nuclei.

## 4. Discussion

As OvCa is one of the most lethal gynaecological malignancies, there is a clear need for robust models that will help uncover the molecular mechanisms underpinning the disease development, growth, metastasis, and even potential therapeutic responses. Cancer modelling over the decades has progressed from crude anatomy to *in vitro* cultures, *in vivo* animal models and now to *in vitro* 3D cultures capable of recapitulating *in vivo* systems and associated TME. In this meta-analysis, we have examined the impact of various scaffolds on the transcriptomic landscape of ovarian cancer cell lines as well as the differences arising from the 3D culture as compared to the classical 2D approaches.

Initial literature survey has pointed out USA as the spearhead of 3D culture research in cancer, covering over 50% of published output in the field. Similar to what is observed in 2D cultures, immortalised cell lines take the forefront with SKOV-3 as the most frequently used option, while primary patient samples are used at a reduced rate. Additional cell lines used are OVCAR3, A2780, PEO1 and OVCAR8. The cell line distribution highlights a strong bias towards White European Ancestry. The percentage of East Asian 3D models in the literature are even lower despite associations with early disease onset in Asian women [29], recapitulating the need for engaging ethnic population in cancer research.

Further analysis shows that the associated subtypes of the cell lines used, align closely with the trend seen in actual global incidence rates of OvCa subtypes. HGSOC is the most frequent of epithelial OvCa subtypes encapsulating 70% of global cases [30], making this subtype a prime dataset to study in this work assessing the variability in 3D culture with respect to classic 2D experiments. It must be noted though, that *in vitro* work requires long-time investment, with relevant models, especially in OvCa, a commodity. With the advance of tissue culture techniques towards more physiological relevant systems however, researchers must strive to use validated and up to date cell lines or note their limitations in disease modelling to maintain reliable and repeatable data.

In this study, we also demonstrate how scaffolds recapitulate the ECM necessary for cell differentiation and the growth of 3D structures [24]. In OvCa modelling, where a 2D counterpart has been used for comparison the most frequent scaffolds utilised by researchers are Matrigel, hanging drop, low attachment plates and hydrogel.

Hanging drop is particularly useful for assessing diffusion gradients in an accessible format [31]. In terms of OvCa this method has been utilised in toxicity screening assays for monitoring chemoresistance in drugs such as cisplatin and Niraparib [18], [32]. Grown in ultra-low attachment plates, OvCa cells show altered mitochondrial function through augmented extracellular acidification rates [33]. Re-sensitisation to treatments in cell lines previously thought resistant are also evident using this method, with a number of BRCA wildtype epithelial OvCa cell lines responding to platinum-based therapeutics and showing an increased rate in apoptosis [34]. Cultures, such as those arising from ovarian malignancies, grown in Matrigel often maintain histological features, genetic profiles, and intra-tumoral heterogeneity, similar to the *in vivo* tumour [35]. Matrigel has also proven an effective model of early-stage angiogenesis in an array of cancers including HGSOC [17]. It must be noted that 3D cultures are often chosen to support the principles of the 3Rs (Replacement, Reduction and Refinement) towards more ethical use of animals [36], [37]. Interestingly, OvCa cell migration, cell communication, and chemotherapeutic response have all been successfully modelled using hydrogel, a plant-based alternative to animal-derivative scaffolds. Here cultures show greater similarity to *in vivo* mouse models and clinical data than that of 2D cultures [37].

### Genetic profile of cells grown in 2D vs. 3D

Leveraging the data from the Gene Expression Omnibus (GEO) and the Sequence Read Achieve allowed us to create a detailed picture of the genomic landscape of ovarian cancer cell lines in 3D cultures using three distinct scaffolds: Matrigel, agarose and collagen. All OvCa cell lines showed a high level of differential regulation with an average of 551 DEGs per data set ranging from 234 DEGs as the minimum and 1429 DEGs as the maximum. The HGSOC cell lines OVCAR8 used across multiple studies allowed us to identify key genes and biological process that are hallmarks of 3D culture as well as potential biomarkers of growth environment for the examined scaffolds. Specifically, our analyses highlight a set of 8 genes, namely DDIT4, ANGPTLA, SELENBP1, SULF1, GAL3ST1, TNFAIP3, LLNLR-263F3.1, MUC12 that show statistically significant differential expression patterns in 3D systems as compared to 2D irrespective of the scaffold used. Furthermore, 13 genes have shown an environment specific expression pattern. The top 16 DEGs between 3D and 2D OVCAR8 were also identified. Of note many of the genes identified are key regulators of inflammation and immune response such as C3, CXCL8 (IL-8), SLPI, CXCL1, CXCL2, ILI beta, IL6, CCL20, IFI27 and CFB [38]–[40]. Furthermore, many of the top genes also show structural importance within the ECM i.e. LAMC2, PI3, FN1, and MMP1. Dysregulation of the matrix metalloproteinase, MMP1, is associated with basement membrane degradation and subsequent peritoneal dissemination in OvCa and is correlated with poor patient prognosis [41]. The remaining DEGs, DDIT4 and ANGPTL4, were recently identified as candidate genes for prediction of survival out comes in lung cancer and OvCa patients [42], [43]. Elevated levels of these glycolysis related genes were also seen to negatively affect progression free survival in patients with OvCa [43].

The functional enrichment scores of OVCAR8 cells grown in Matrigel, agarose and collagen, compared with standard 2D mono-layer controls presented a unique expression profile with close relation seen between the 2D samples. However, the 3D collagen OVCAR8 cells expressed a higher degree in variability compared with the other 3D OVCAR8’s which show comparatively similar profiles. Earlier studies have suggested that this model is more similar to the *in vivo* environment as it captures 3D growth alongside omental fibroblasts [44].

The top biological processes associated with the DEGs identified between the 2D and 3D include glycolysis, KRAS signalling, coagulation, TNF alpha signalling via NF-κB, complement and inflammatory response. These processes are frequently altered in cancer and are often difficult to model in 2D systems [45]. Glycolysis in particular is often augmented in cancer cells with increased utilisation of this pathway indicative of the Warburg effect [46]. Similar metabolic changes are also evident in 3D colorectal cancer cells when compared to 2D [47]. The inclusion of these processes in the data verifies numerous studies where 3D cells are shown to express more biological relevance to *in vivo* systems than 2D cell cultures, through the expression of pathways typically associated with *in vivo* environments [45], [47]–[51].

Furthermore, some cancer related hallmarks were also highlighted as differentially regulated in the 3D OvCa cells when compared with the 2D samples. Hallmarks of particular interest include apoptosis, oxidative phosphorylation, MYC pathways, ROS, EMT, KRAS signalling, angiogenesis and hypoxia. Numerous studies show that the 3D environment influences these key cancer pathways [45], [47]; here we show that regardless of scaffold the processes are still heavily influenced when grown in 3D. Apoptosis, EMT, KRAS signalling and hypoxia as well as angiogenesis were some of the key cancer associated processes enhanced in 3D growth. Additional processes included complement and inflammatory response pathways which are important factors of tumour immune evasion. Another pathway often seen in cancers was IL6-JAK-STAT3, which is a proliferative driver often implicit with OvCa angiogenesis and tumour metastasis [52].

Moreover, based on the expression profile of OVCAR8 cells grown in 3D vs. 2D, we identified a panel of genes specific to OVCAR8 when grown in different gel-based scaffolds using Omics Playground importance score ranking [27]. The expression profile of these genes was unique to the specific scaffold when compared with the 2D OVCAR8. Biomarkers specific to OvCa cells grown in agarose compared with 2D include: C3, MMP1, ILIB and CCL20. The three biomarkers identified for collagen include: IFI27, COL3A1 and IFI27. Matrigel however only showed one unique marker, DDIT4 a stress included regulator of mTOR previously mentioned for its association with progression free survival in OvCa [43]. Future work should explore the relevance of these markers and the influence they hold within the OvCa TME.

Next, we explored the impact of cell line on various scaffolds and showed that there is a close relationship between the two suggesting that in order to recover the tissue specific behaviour in a model 3D culture, a lot of care must be given to the choice of cell line and scaffold, in order to remove potential experimental biases. Furthermore, the condition specific gene expression patterns suggested that a number of genes can be used as environment biomarkers.

Finally, we explored the impact of transcriptional changes in real time by looking at phenotypic changes of cells grown in 3D vs 2D cultures. Our experiment have shown that SKOV-3 cells grown in hydrogel are clustering in the simple spheroids, precursors of higher order organoid formations.

In summary this meta-analysis assessed the current landscape of 3D OvCa within the literature and provided a complex expression profile of OvCa cells grown in 3D. Our transversal comparison of various scaffolds allowed us to highlight the variability that can be induced by various scaffolds in the transcriptional landscape as well as identifying key genes and biological processes that are hallmarks of cancer cells grown in 3D cultures. Moreover, the identification of growth environment biomarkers will allow us to monitor in the future the suitability of 3D culture to recapitulate tissue complexity.

## Supplementary Materials

The following supporting information is available: Figure S1: Top Enriched Gene sets for 2D vs. 3D OVCAR8; Table S1: Cell line information and associated accession codes.

## Author Contributions

For research articles with several authors, a short paragraph specifying their individual contributions must be provided. The following statements should be used “Conceptualization, C.S and R.K.; methodology, C.S., R.K., S.P., EK and B.B.; software, C.S. and B.B.; validation, R.K., S.P. and C.S.; formal analysis, R.K., BB; investigation, R.K., E.K., C.S.; resources, C.S., I.K., H.R., J.H., and M.H.; data curation, R.K. and C.S.; writing—original draft preparation, R.K.; writing—review and editing, R.K., C.S., and E.K.; visualization, C.S.; supervision, E.K., I.K., M.H.; project administration, E.K., H.R., J.H., funding acquisition, I.K., H.R., M.H. All authors have read and agreed to the published version of the manuscript

## Funding

This study was funded through the Cancer Treatment & Research Trust and University Hospitals Coventry and Warwickshire NHS Trust (grant no. 12899).

## Data Availability Statement

RNAseq and Array data can be found via the following NCBI accession codes: PRJNA472611, PRJNA530150, PRJNA564843, PRJNA564843, PRJNA232817, PRJNA318768. A full list of samples can be viewed in Supplementary table 1.

## Acknowledgments

The authors acknowledge the Biocenter Finland (BF) and Tampere Imaging Facility (TIF) for the service.

## Conflicts of Interest

The authors declare no conflict of interest.

## Appendix A

**Supplementary Table 1.** Cell line information and associated accession codes

| Accession Number | SRA code | Cell line | Subtype | Condition | Scaffold |
| --- | --- | --- | --- | --- | --- |
| PRJNA472611 | SRR7204219 | A1847 | Carcinoma | 3D | Agarose |
|  | SRR7204220 | A1847 | Carcinoma | 2D | / |
|  | SRR7204221 | A2780 | HGSOC | 3D | Agarose |
|  | SRR7204222 | A2780 | HGSOC | 2D | / |
|  | SRR7204223 | OVCAR3 | HGSOC | 3D | Agarose |
|  | SRR7204224 | OVCAR3 | HGSOC | 2D | / |
|  | SRR7204225 | OVCAR4 | HGSOC | 3D | Agarose |
|  | SRR7204226 | OVCAR4 | HGSOC | 2D | / |
|  | SRR7204227 | OVCAR5 | HGSOC | 3D | Agarose |
|  | SRR7204228 | OVCAR5 | HGSOC | 2D | / |
|  | SRR7204231 | OVCAR10 | HGSOC | 3D | Agarose |
|  | SRR7204232 | OVCAR10 | HGSOC | 2D | / |
|  | SRR7204233 | OVCAR8 | HGSOC | 3D | Agarose |
|  | SRR7204234 | OVCAR8 | HGSOC | 2D | / |
|  | SRR7204235 | SKOV-3 | HGSOC | 3D | Agarose |
|  | SRR7204236 | SKOV-3 | HGSOC | 2D | / |
|  | SRR7204237 | PEO1 | HGSOC | 3D | Agarose |
|  | SRR7204238 | PEO1 | HGSOC | 2D | / |
|  | SRR7204229 | C30 | Carcinoma | 3D | Agarose |
|  | SRR7204230 | C30 | Carcinoma | 2D | / |
|  | SRR7204242 | C70 | Carcinoma | 3D | Agarose |
|  | SRR7204241 | C70 | Carcinoma | 2D | / |
|  | SRR7204240 | UPN275 | MAC | 3D | Agarose |
|  | SRR7204239 | UPN275 | MAC | 2D | / |
| PRJNA530150 | SRR8823257 | OVCAR8 | HGSOC | 2D | / |
|  | SRR8823258 | OVCAR8 | HGSOC | 2D | / |
|  | SRR8823259 | OVCAR8 | HGSOC | 2D | / |
|  | SRR8823260 | OVCAR8 | HGSOC | 2D | / |
|  | SRR8823265 | OVCAR8 | HGSOC | 3D | Matrigel |
|  | SRR8823266 | OVCAR8 | HGSOC | 3D | Matrigel |
|  | SRR8823267 | OVCAR8 | HGSOC | 3D | Matrigel |
|  | SRR8823268 | OVCAR8 | HGSOC | 3D | Matrigel |
|  | SRR8823273 | OVCAR8 | HGSOC | 2D | / |
|  | SRR8823273 | OVCAR8 | HGSOC | 2D | / |
|  | SRR8823273 | OVCAR8 | HGSOC | 2D | / |
|  | SRR8823273 | OVCAR8 | HGSOC | 2D | / |
|  | SRR8823280 | OVCAR8 | HGSOC | 3D | Matrigel |
|  | SRR8823281 | OVCAR8 | HGSOC | 3D | Matrigel |
|  | SRR8823282 | OVCAR8 | HGSOC | 3D | Matrigel |
|  | SRR8823283 | OVCAR8 | HGSOC | 3D | Matrigel |
| PRJNA564843 | SRR10096845 | OVCAR8 | HGSOC | 2D | / |
|  | SRR10096844 | OVCAR8 | HGSOC | 2D | / |
|  | SRR10096843 | OVCAR8 | HGSOC | 2D | / |
|  | SRR10096841 | OVCAR8 | HGSOC | 3D | Collagen |
|  | SRR10096842 | OVCAR8 | HGSOC | 3D | Collagen |
|  | SRR10096840 | OVCAR8 | HGSOC | 3D | Collagen |
|  | SRR10096839 | OVCAR4 | HGSOC | 2D | / |
|  | SRR10096838 | OVCAR4 | HGSOC | 2D | / |
|  | SRR10096837 | OVCAR4 | HGSOC | 2D | / |
|  | SRR10096836 | OVCAR4 | HGSOC | 3D | Collagen |
|  | SRR10096835 | OVCAR4 | HGSOC | 3D | Collagen |
|  | SRR10096834 | OVCAR4 | HGSOC | 3D | Collagen |
|  | SRR10096828 | Kuramochi | HGSOC | 3D | Collagen |
|  | SRR10096829 | Kuramochi | HGSOC | 3D | Collagen |
|  | SRR10096830 | Kuramochi | HGSOC | 3D | Collagen |
|  | SRR10096831 | Kuramochi | HGSOC | 2D | / |
|  | SRR10096832 | Kuramochi | HGSOC | 2D | / |
|  | SRR10096833 | Kuramochi | HGSOC | 2D | / |
| PRJNA232817 | GSM1300206 | IGROV-1 | EAC | 2D | / |
|  | GSM1300207 | IGROV-1 | EAC | 2D | / |
|  | GSM1300208 | IGROV-1 | EAC | 2D | / |
|  | GSM1300209 | IGROV-1 | EAC | 3D | Low Attachment |
|  | GSM1300210 | IGROV-1 | EAC | 3D | Low Attachment |
|  | GSM1300211 | IGROV-1 | EAC | 3D | Low Attachment |
| PRJNA318768 | GSM2125384 | HEY | HGSOC | 2D | / |
|  | GSM2125385 | HEY | HGSOC | 2D | / |
|  | GSM2125386 | HEY | HGSOC | 2D | / |
|  | GSM2125387 | HEY | HGSOC | 2D | / |
|  | GSM2125388 | HEY | HGSOC | 3D | Hanging drop |
|  | GSM2125389 | HEY | HGSOC | 3D | Hanging drop |
|  | GSM2125390 | HEY | HGSOC | 3D | Hanging drop |
|  | GSM2125391 | HEY | HGSOC | 3D | Hanging drop |

Further gene set enrichment analysis revealed that when grown in 3D many processes associated with hallmarks of cancer were also differentially regulated. Of note key processes that often show enhanced presentation in 3D growth such as angiogenesis, apoptosis and hypoxia all exhibit enrichment.

**Supplementary Figure 1.**
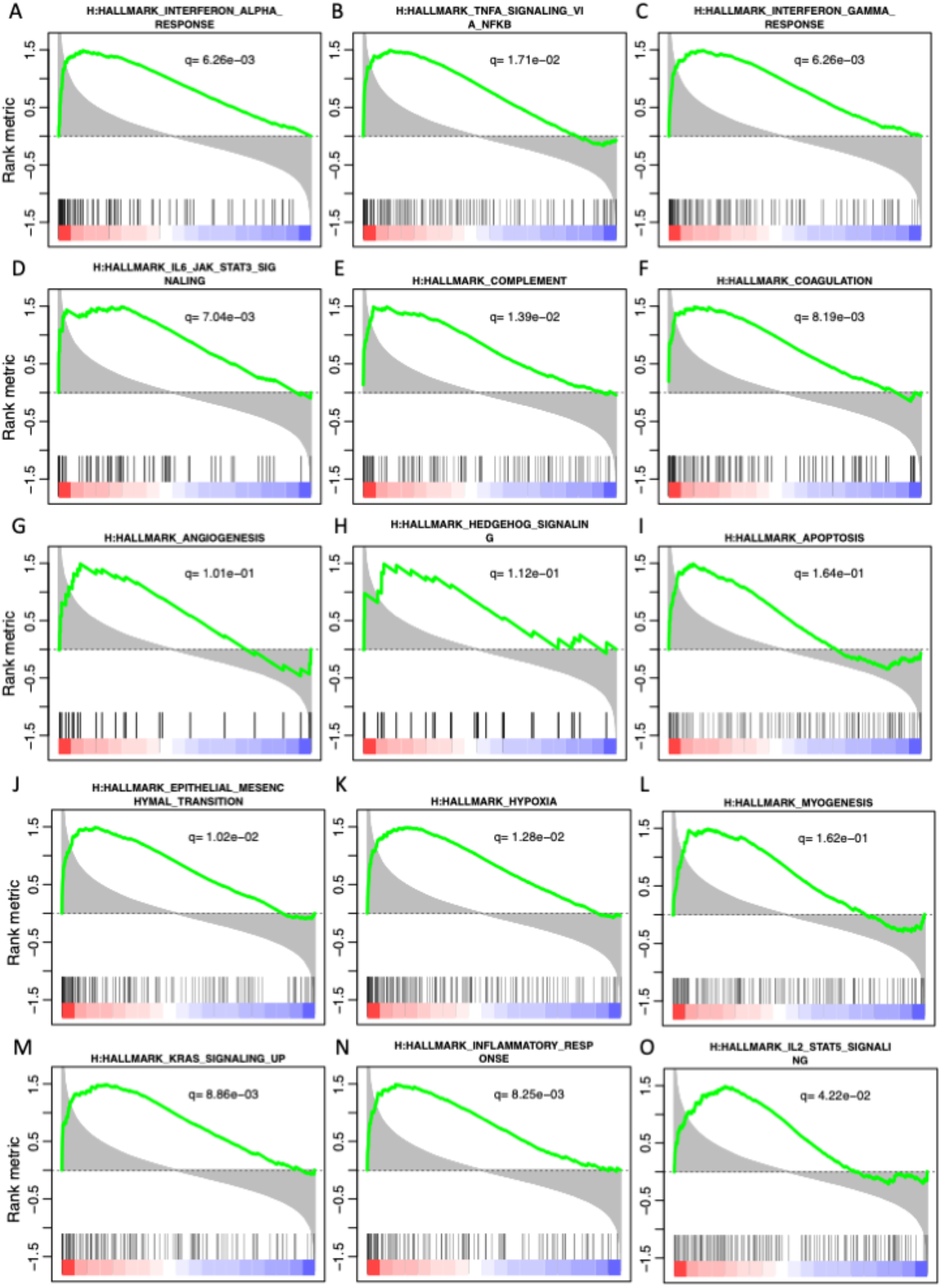
Top Enriched Gene sets for 2D vs. 3D OVCAR8. Combined panel of enrichment curves showing processes associated with cancer hallmarks. **(A)**, Interferon Alpha Response; **(B)**, TNF-a signalling; **(C)**, Interferon Gamma Response; **(D),** IL-6-JAK-STAT3 signalling; **(E)**, Complement; **(F)**, Coagulation; **(G)**, Angiogenesis; **(H)**, Hedgehog signalling; **(I)**, Apoptosis; **(J),** Epithelial Mesenchymal Transition; **(K)**, Hypoxia; **(L)**, Myogenesis; **(M)**, KRAS signalling; **(N)**, Inflammatory Response; **(O)**, IL-2 STAT3 signalling and. Black vertical bars represent gene rank using shorted list metric. Green curve corresponds to “running statistics” of the enrichment score (ES).

## Notes

### Competing Interest Statement

The authors have declared no competing interest.

